# A key role of astrocytic calcium dynamics to link neuronal activity with the BOLD signal

**DOI:** 10.1101/2021.04.23.441146

**Authors:** Federico Tesler, Marja-Leena Linne, Alain Destexhe

**Affiliations:** Paris-Saclay University, CNRS, Paris-Saclay Institute of Neuroscience (NeuroPSI), 91198 Gif-sur-Yvette, France; Tampere University, Faculty of Medicine and Health Technology, 33720 Tampere, Finland

**Keywords:** fMRI, neurovascular coupling, astrocytes, calcium signaling

## Abstract

Functional magnetic resonance imaging (fMRI) relies on the coupling between neuronal and vascular activity, but the mechanisms behind this coupling are still under discussion. Recent experimental evidence suggests that calcium signaling may play a significant role in neurovascular coupling. However, it is still controversial where this calcium signal is located (in neurons or elsewhere), how it operates and how relevant is its role. In this paper we introduce a biologically plausible model of the neurovascular coupling and we show that calcium signaling in astrocytes can explain the main aspects of the dynamics of the coupling. We find that calcium signaling can explain so-far unrelated features such as the linear and non-linear regimes, the negative vascular response (undershoot) and the emergence of a (calcium-driven) Hemodynamic Response Function. These features are reproduced here for the first time by a single model of the detailed neuronal-astrocyte-vascular pathway. Furthermore, we analyze how information is coded and transmitted from the neuronal to the vascular system and we find that frequency modulation of astrocytic calcium dynamics plays a key role in this process. Finally, our work provides a framework to link neuronal activity to the BOLD signal, and vice-versa, where neuronal activity can be inferred from the BOLD signal. This opens new ways to link known alterations of astrocytic calcium signaling in neurodegenerative diseases (e.g. Alzheimer’s and Parkinson’s diseases) with detectable changes in neurovascular coupling.

## 1. Introduction

Functional magnetic resonance imaging (fMRI) has become one of the leading neuroimaging techniques in neuroscience. fMRI methods rely on the coupling between local neuronal activity and cerebral blood flow (CBF) [1, 2]. In normal conditions an increase in neuronal activity is followed by an increase in CBF, which supplies the necessary oxygen and glucose to sustain the cerebral metabolism, a process known as *functional hyperaemia*. In a healthy brain, the increase in oxygen supply overpasses the oxygen demand generating an increase in the local level of blood oxygenation, which is captured by the BOLD (Blood-Oxygen-Level-Dependent) signal [3]. Although the neurovascular coupling has been extensively demonstrated, the mechanisms behind it remain under discussion.

In recent years it has been shown that BOLD signal has a stronger correlation with synaptic activity than with single neuron or multi-unit spiking activity [2, 4, 5]. In particular glutamatergic synapses are believed to play a central role in CBF activation, presumably by interacting with glutamate receptors on astrocytes and neurons which triggers the production and release of a variety of vasomodulators [6, 7]. Among these vasomodulators we find potassium (K^+^) and hydrogen (H^+^) ions, prostaglandins (PGs), epoxyeicosatrienoic acid (EET) and nitric oxide (NO) [2, 7, 8]. The action of vasodilators on smooth cells of nearby arterioles generates an increase in their volume which leads to an increase in cerebral blood flow. The relative relevance of the different vasodilators (and their molecular pathways) is still under study and is believed to vary among brain regions and cell types [2, 6, 7].

Despite the different sources and effects on the vasculature, one feature that the different coupling mechanisms seem to share is that they are usually associated with some type of intracellular calcium (Ca^2+^) activity. This relation between calcium signaling in the brain and the vascular response has started to be further explored in experiments during the last few years. Calcium activity in neuronal dendrites, whole-cells and astrocytes has been analyzed in relation with the vascular response [6, 9, 10, 11, 12].

There is however little understanding on how the processing of information is performed by the calcium signal and how it operates on the formation of the BOLD signal. To understand the role of calcium signaling in the generation of the BOLD response is the goal of this work. In particular, we will focus on calcium signaling in astrocytes and we will provide evidence, via modelling and simulations, of the central role that it can be playing in the coupling.

To perform our study we introduce and analyze a biologically plausible model of the neurovascular coupling which incorporates recent experimental findings and modelling tools at the three levels of the system (neuronal, astrocytic, vascular). For concreteness we adopt the pathway defined by the calcium activity in astrocytes as the main mediator between neuronal activity and the vascular system. We will consider the release of the prostaglandin PGE2 by astrocytes as the principal vasomodulator acting in the coupling. Neuronal (and synaptic) activity will be modeled via a recently developed mean-field description of a network of Adaptive Exponential(AdEx) integrate-and-fire neurons [13], which allows us to explicitly integrate neuronal adaptation.

Starting from a relatively detailed description we will focus on fundamental aspects of the fMRI phenomenology (such as main experimental protocols, the Hemodynamic Response Function, the linearity of the coupling, the post-stimulus undershoot and the effects of neuronal adaptation) and we will analyze the role of the calcium signal in these processes. Previous studies with models of the neuro-astro-vascular interaction exists [14, 15, 16], but to the best of our knowledge, this is the first work that accounts for the above experimental observation of fMRI.

We show that information transmission by the calcium signal operates mainly via frequency coding with a small contribution of amplitude modulation. We also show that in our model calcium signaling is responsible for both linear and non-linear aspects of the neurovascular coupling. In section 3.3 we show that a calcium-driven Hemodynamic Response Function (HRF) can be generated and that it is equivalent to the one observed experimentally. We corroborate this by comparing our results with the canonical HRF and we reproduce our simulations using the HRF formalism. We also show that the calcium activity is a better predictor of the BOLD response within this context. In section 3.4 we study situations where the BOLD signals can become negative (post-stimulus undershoot), a well-know feature observed in fMRI experiments. We show that an undershoot in the calcium activity can cause an undershoot in the BOLD signal via a decrease in CBF. Finally, in section 3.5 we study the effects of neuronal adaptation on the calcium and BOLD signals. We show that the calcium signal can capture the effects of neuronal adaptation via calcium-spike frequency adaptation and transmit it to the BOLD signal. In particular we show that adaptation may play a major role in the post-stimulus undershoot, which further stresses the usefulness of adopting a mean-field formalism where adaptation is present. We end by showing how information about the incoming and recurrent neuronal activity can be extracted from the BOLD signal in our modeling framework.

## 2. The model

The model of the neurovascular coupling that we propose consists in a feedforward system comprising the neuronal activity, the calcium dynamics in astrocytes and the vascular response. A diagram of the model is shown in Fig. 1. Neuronal activity will be described via a mean-field model of a network of AdEx neurons. Calcium dynamics in astrocytes will be described via the traditional Li-Rinzel model with recent improvements by De Pittà et al. (2009) to incorporate IP3 dynamics by glutamate activation [17, 18]. To describe the communication between astrocytes and the vascular system we will model the production of the prostaglandin PGE2 (via the arachidonic acid (AA) cascade) and the action of this vasomodulator on arteriolar smooth-cells which induces the changes in cerebral blood flow. Finally, the BOLD signal generation is modeled via the Balloon model [19]. The details of the model are introduced in the following.

**Figure 1:**
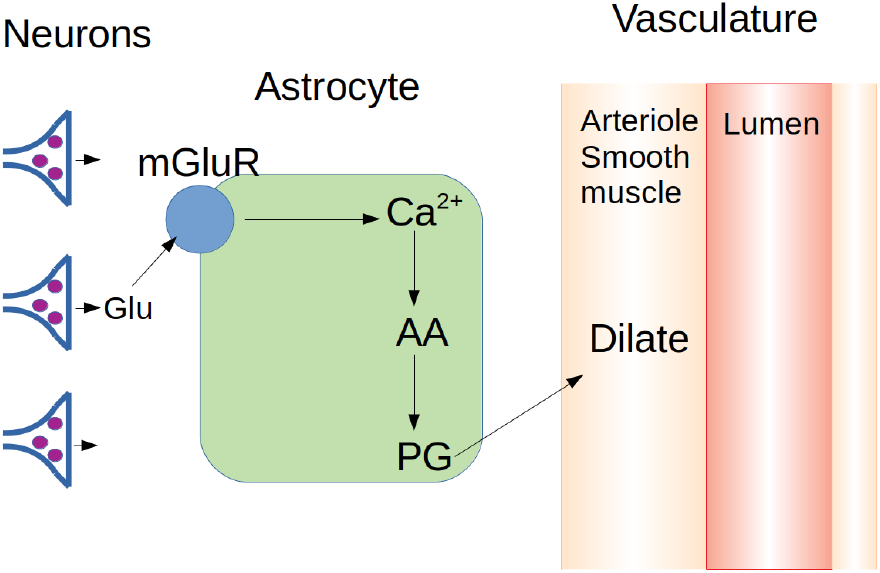
Diagram of the model of neurovascular coupling. We consider a feedforward system that starts at the neuronal level. Glutamate spilled out from synaptic clefts toward the perisynaptic space activates, through metabotropic glutamate receptors (mGluRs), the calcium (Ca^2+^) signaling cascade in astrocytes. The increase in cytosolic Ca^2+^ levels in astrocytes induces the production and release of vasomodulators (prostaglandin, PG) derived from free arachidonic acid, leading to the variations of the arteriole volume.

### 2.1. Mean-field neuronal model and synaptic release

To model the neuronal activity we will consider a network of adaptive exponential-integrate-and-fire neurons (AdEx) [20]. We will consider a directed network made by 10000 AdEx neurons, with 80% of excitatory and 20% of inhibitory neurons. Neurons in the network are randomly connected with probability *p* = 5% and interact via conductance based synapses (see Supp. Inf.Materials and Methods for details). We will focus on the mean activity of the population. The mean-field equations for the AdEx network are given to a first-order by [13]:

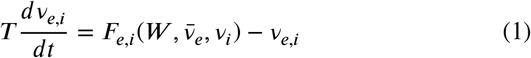

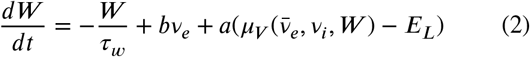

where *v*_*e,i*_ is the mean neuronal firing rate of the excitatory and inhibitory population respectively, *W* is the mean value of the adaptation variable, *F* is the neuron transfer function (i.e. output firing rate of a neuron when receiving excitatory and inhibitory inputs with mean rates *v*_*e*_ and *v*_*i*_ and with a level of adaptation W), *a* and *b* are the sub-threshold and spiking adaptation constants, *t*_*w*_ is the characteristic time of the adaptation variable and *T* is a characteristic time for neuronal response (we adopt *t*_*w*_ = 1 s and *T* = 5 ms). Values for all the parameters are given in Supp. Inf.Table 1. The stimulation of the network is simulated by an (excitatory) external drive *v*_*ext*_, which represents the input from a nearby neuronal populatio n and so 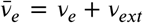. The network exhibits no spontaneous activity for the parameters used. Derivation of the mean field equations can be found in Ref. [13].

**Table 1:** Model parameters for equations described in the main text. ^*^: This value of *g*_*r*_ incorporates the cleft/perisynaptic volume ratio and the percentage of glutamate spilled to the perysinaptic space.

| **AdEx Mean Field Model** |  |  |
| --- | --- | --- |
| $N_E$ | #excit. neurons | 8000 |
| $N_I$ | #inhibit. neurons | 2000 |
| $p$ | Connection probability | 5% |
| $T$ | Neuronal characteristic time | 5ms |
| $\tau_W$ | Adaptation time constant | 1s |
| $b_E$ | Adaptation spike-triggered constant (excit. neurons) | 60pA |
| $b_I$ | Adaptation spike-triggered constant (inhibit. neurons) | 0 |
| $a$ | Adaptation sub-threshold constant (excit/inhibit neurons) | 0 |
| $E_L$ | Leakage reverse potential (excit/inhibit neurons) | -70mV |
| $g_r^*$ | Constant of glutamate release by recurrent activity | $4.2 \times 10^{-3} \text{mM/S}$ |
| **Astrocyte** |  |  |
| $\tau_{AA}$ | Relaxation time AA | 1s |
| $O_{AA}$ | Max. rate of AA production | $5 \times 10^{-3} \text{mMs}^{-1}$ |
| $K_{AA}$ | Michaelis constant AA | $0.8 \text{e-}3 \times 10^{-3} \text{mM}$ |
| $\tau_{PG}$ | Relaxation time PGE2 | 1s |
| $O_{PG}$ | Max. rate of PG production | $2 \times 10^{-3} \text{mMs}^{-1}$ |
| $K_{PG}$ | Michaelis constant PGE2 | $1 \text{e-}3 \times 10^{-3} \text{mM}$ |
| **Vascular System** |  |  |
| $\tau_R$ | Relaxation time PGE2 receptor | 0.8s |
| $O_R$ | PGE2 binding rate | $0.2 \times 10^4 (\text{mMs})^{-1}$ |
| $\tau_{cAMP}$ | Relaxation time *cAMP* | 2s |
| $O_{cAMP}$ | Max. rate of *cAMP* production | $2 \times 10^{-3} \text{mMs}^{-1}$ |
| $D_A$ | Max. Arteriole dilation | 5 |
| $K_{VA}$ | Arteriole-*cAMP* half dilation | $5 \times 10^{-3} \text{mM}$ |

Synaptic glutamate release can be straightforwardly estimated from the mean-field model. Glutamate concentration ([Glu]) is proportional to the mean conductance of excitatory synapses (*s*) obtained from the (conductance-based) meanfield model [13], *s* = *v*_*e*_*pN*_*e*_*Q*_*e*_ *τ* _*e*_, where *N*_*e*_ is the number of excitatory neurons, *Q*_*e*_ is the excitatory quantal conductance (conductance change induced by a single spike) and *τ* _*e*_ is the conductance decay time (see Supp. Inf. for details and Supp. Inf. Table 1 for the corresponding parameter values). Thus, [Glu]=*gs*, where *g* is a proportionality constant. For a better analysis, we will separate the contribution of the recurrent excitatory synaptic activity from the one of the external excitatory input. Thus, the mean glutamate concentration in the synaptic cleft is given by:

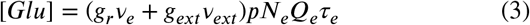

where *g*_*r/ext*_ is the proportionality constant accounting for the contribution to glutamate release of the recurrent and external excitatory synaptic activity respectively.

### 2.2. Astrocyte calcium dynamics and arachidonic acid cascade

In this section we present the model used to describe the intracellular calcium dynamics and vasomodulator production by astrocytes. We assume that part of the neuronal glutamate released to the synaptic cleft spills out toward the perisynaptic space and activates metabotropic glutamate receptors (mGluRs) of nearby astrocytes. The activation of mGluRs induces an increase of intracellular cytosolic inositol trisphosphate (IP3) that triggers the release of calcium (Ca^2+^) from the endoplasmic reticulum (ER) towards the cytosol. This generates a transient increase in cytosolic Ca^2+^ which is later reabsorbed by the ER. To model this glutamate-induced calcium dynamics in the astrocyte we adopt a recent version of the Li-Rinzel model [17] which incorporates a detailed dynamics of *Ca*^2+^-dependent production and degradation of IP3 [18, 21]. For details on the model see Refs.[18, 17, 21] and Supp. Inf. Materials and Methods.

The resultant increase in cytosolic Ca^2+^ activates phospholipase A2, which induces the production of arachidonic acid (AA) from membrane phospholipids. This in turn leads to the production and release of vasomodulators derived from AA, in particular the vasodilator PGE2 [2, 7, 8]. The production of AA and PGE2 will be modeled via Michaelis-Menten type of equations [14]:

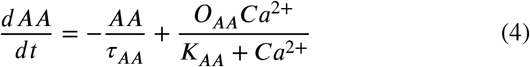

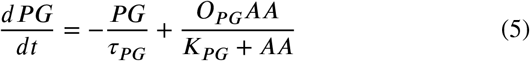

where *τ*_*AA*_ and *τ*_*PG*_ are the time constants for AA and PGE2 respectively, *K*_*PG*_ and *K*_*AA*_ are the Michaelis constants and *O*_*PG*_, *O*_*cAMP*_ are the maximum rate of production of AA and PGE2 respectively. We note that this model describes a single cell activity. Vascular activation is most likely not carried out by a single astrocyte (or single astrocyte process) but by the many astrocytes linked with the local vasculature and neuronal population. Nevertheless, it has been seen that *in vivo* the activity of astrocytes tend to be restricted to independent single-cell activity [22, 23, 24, 25] (rather than intercellular calcium waves as seen in *in vitro* experiments [26]). Thus, we will adopt this single-cell model as a representative unit of the astrocytes associated with the same neuronal population.

### 2.3. Vascular response and BOLD signal

We will assume that the vascular response starts at the arteriole level. In our model, PGE2 released from astrocytes binds to EP4 prostaglandin receptors on smooth cells of arterioles inducing an increase of cyclic adenosine monophosphate (cAMP) concentration in the smooth cells. This generates the activation of protein kinase A and a decrease in the phosphorylation of the myosin light chain. Contraction of arterioles has been proposed to be proportional to the concentration of phosphorylated myosin [15]. Following this line, we will assume that arteriole dilation is proportional to the concentration of cAMP. The equations describing the fraction of activated PGE2 receptors and the cAMP production in arterioles are:

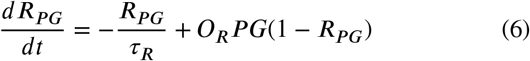

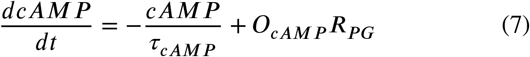

where *R*_*PG*_ is the portion of activated PGE2 receptors, *PG* is the concentration of PGE2, *τ*_*cAMP*_ is the relaxation time for cAMP, and *O*_*R*_, *O*_*cAMP*_ are constants. Normalized relative arteriole volume is given by:

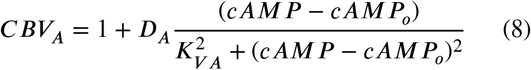

where *CBV*_*A*_ is the normalized change of arteriolar cerebral blood volume relative to its basal value, *cAMP*_*O*_ is the basal concentration of cAMP and *D*_*A*_,*K*_*VA*_ are constants.

The relation between CBV and CBF is usually assumed to follow a power law. The value of the exponent has been observed to depend on the specific brain region and vasculature level [27, 28]. In our model the relative change in cerebral blood flow evoked by changes in *CBV*_*A*_ is given by:

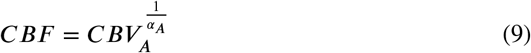

where *α*_*A*_ is a constant (a different power *α*_*V*_ is used for the venule blood volume *CBV*_*V*_, see Supp. Inf. Materials and Methods).

Finally, to generate the BOLD signal evoked by the changes of CBF we will adopt the well-known Balloon model [19]. This model describes the change in dexohemoglobin concentration (which originates the BOLD signal) generated by an increase in CBF. A rough approximation of the BOLD signal obtained from the Balloon model in terms of CBF is given by [29]:

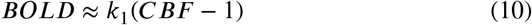

where *k*_1_ is a constant. This approximation provides a good intuition of the expected BOLD response and is given here for reference, however it fails to reproduce some key features such as the post-stimulus undershoot related with slow venule volume relaxation (analyzed later in the paper). For all the simulations presented in this paper we use the full Balloon model as presented in the Supp. Inf. (Materials and Methods). For details on the Balloon model see Refs. [19, 30] and Supp. Inf. Materials and Methods.

## 3. Results

In the following we present the results obtained from our model. First we analyze the time course of the main variables in the vascular pathway and we reproduce some key experimental results from fMRI measurements. In particular, we show that our simulations can correctly reproduce the results of the two main experimental protocols used in BOLD imaging, namely i) event-related protocols and ii) block protocols. Then we proceed with the study of information coding and transmission in our system (3.2); the analysis of the linear coupling and the Hemodynamic Response Function formalism (3.3); the analysis of the undershoot of the BOLD signal in the context of the calcium pathway (3.4); and the effects of neuronal adaptation on the calcium signal and the BOLD response (3.5).

### 3.1. Event-related and block protocols

In this section we show how the calcium pathway for neuro-vascular coupling operates and we show that we can correctly reproduce the results from the main experimental protocols used in fMRI, which provides a first validation for our model.

Experimental protocols used for BOLD imaging are usually divided in two groups [1, 31, 32]: i) event-related protocols, where short single stimulus are presented with long inter-stimulus intervals (tens of seconds, equal or larger than typical BOLD response time-course); and ii) block protocols, where multiple stimuli are presented with very short inter-stimulus intervals, followed by a longer ‘inter-block’ interval after which a new set (‘block’) of stimuli is presented. In both protocols stimulus duration can be in the order of 4s or less. Variations and mixed-type protocols are also used. For a detailed analysis of the protocols and their advantages see Refs. [1, 31, 32, 33, 34]. We notice that these two protocols are related to fundamental features of the neurovascular coupling, which are the vascular response to an impulse stimulus and the response to a superposition of stimulus. The analysis of of these two features constitutes an important part of this work.

The results of our simulations for the two protocols are shown in Fig. 2. Comparison with experimental results are shown in panel (b). Except otherwise indicated, the stimulus used in the paper will consist in a pulse of the external input *v*_*ext*_ as shown in the top panel of Fig.2.a (the width of the pulse corresponds to the duration of the stimulus). The response obtained for a single (short) event is usually known as the Hemodynamic Response Function (HRF) and correspond to the minimum BOLD response in functional hyperemia. The same response is obtained for any stimulus of shorter duration surpassing a certain threshold in time and intensity. In our model, this minimum response corresponds to the activation of a calcium-spike as we will show later. The HRF is analyzed in detail in section 3.3. From the simulation of the block protocol we see that we can correctly capture the response to the superposition of events (further analysis on this is developed in section 3.3).

**Figure 2:**
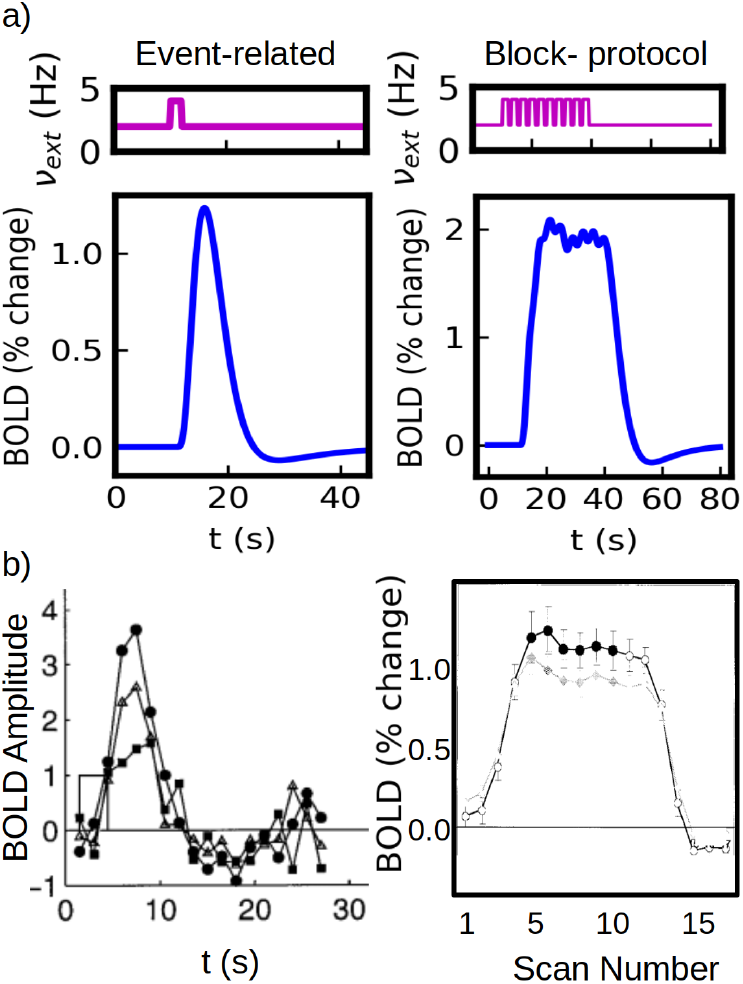
Standard experimental protocols for fMRI measurements. a) Simulations of the event-related and block protocols. The event-related case consists in the application of a single stimulus of 2s. The block-protocol consists in the application of 10 stimuli of 2 seconds each, separated by an inter-stimulus interval of 1 second. b) Experimental results for the two standard protocols. Left: results from an event-related protocol for three different stimulus strength (stimulus duration of 3 seconds), adapted from Ref. [35]. Right: results from a block protocol with 2.5 seconds stimulus and an inter-stimulus interval of 0.5 seconds, adapted from Ref. [34].

To provide a better understanding of the mechanism of the BOLD generation in our system we show in Fig. 3 the time course of the main variables of the model for stimuli of two different durations (4s and 24s). In Fig.3 (top panels) we see that a fast neuronal response is generated after the application of the stimulus, which exhibits a marked overshoot given by the strong adaptation effect. This neuronal response generates an increase in the perisynaptic glutamate concentration (second panels from top) that triggers the activation of the calcium dynamics in the astrocyte (characterized by calcium-spikes as seen in Fig. 3 third panels from top). The Ca^2+^ activity leads to the increase in cerebral blood flow (CBF_*IN*_, bottom panels) driven by the release of the vaso-modulator PGE2 and the increase in arteriole volume (not shown, see Supp. Inf. Materials and Methods for the time course of the remaining variables). We notice that CBF_*IN*_ is linked to the arteriole volume CBV_*A*_. Timing and size differences between arteriole and venule volume changes are also correctly captured by our simulations and are shown in the Supp. Inf. Materials and Methods. As we will see in the next section, the information about the external stimulus is coded in terms of the frequency and amplitude of calcium spikes.

**Figure 3:**
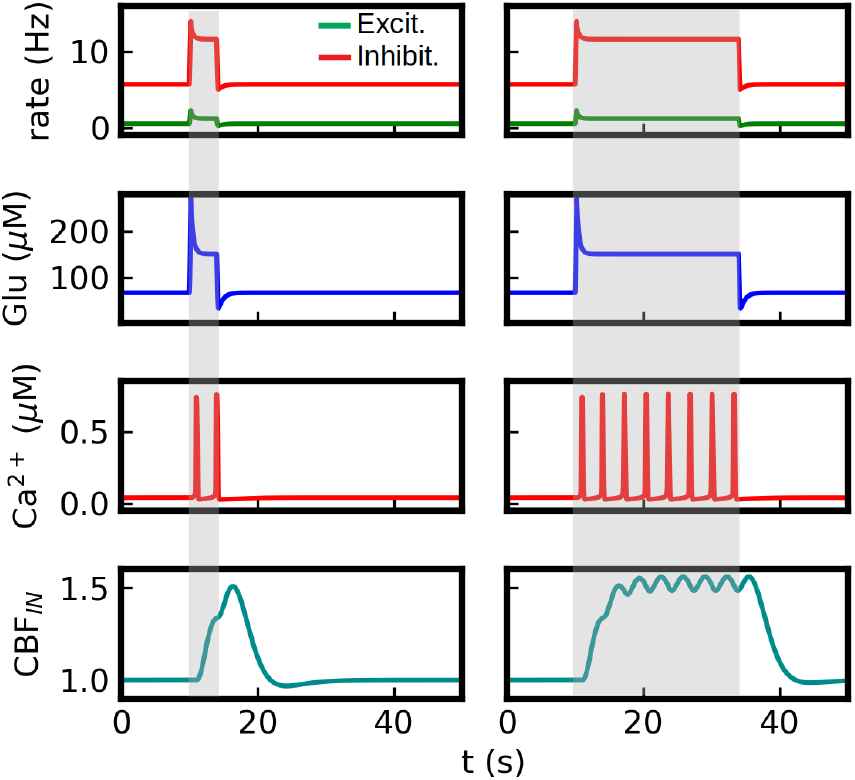
Time course of main model variables for a stimulus of 4 seconds (left) and 24 seconds (right). The interval of stimulus application is indicated by the grey area. The figure illustrates the calcium pathway of neurovascular coupling in the model. The stimulus (represented as an increase in the external input *v*_*ext*_, not shown) generates an increase in neuronal activity (top panel) which induces a rise in the level of perisynaptic glutamate concentration (second panel). Glutamate triggers the activation of calcium dynamics in astrocytes which is characterized by brief calcium ‘spikes’ as seen in the figure (third panels from the top). This leads to the production and release of the vasomodulator PGE2 (not shown, see Supp. Inf. Materials and Methods) which dilates the nearby arteriole and increases the flow of blood into the tissue (bottom panel). The simulations correspond to *v*_*ext*_=4Hz.

In the simulations presented in this section we set *g*_*ext*_ = 0 in Eq.3, meaning that the glutamate release is proportional to the local excitatory activity *v*_*e*_. Nevertheless, we notice that, for the level of neuronal adaptation that we use, the asymptotic activity of the population is mainly driven by the external input with a small contribution of the recurrent activity (and *v*_*e*_ is nearly proportional to *v*_*ext*_, as seen in the next section). The recurrent activity is only relevant during a short period after the stimulus onset (neuronal overshoot), defined by the characteristic time of the adaptation variable (*t*_*w*_). As this characteristic time is short compared to the calcium response, then the BOLD response in these simulations is mainly driven by the asymptotic neuronal activity (and thus by the external input). Thus, the results presented here for the BOLD signal are nearly independent of the *g*_*r/ext*_ values we use. We will use *g*_*ext*_ = 0 through the paper, except otherwise indicated. As we will see in section 3.5, when *t*_*w*_ is comparable to the time of the calcium response, then the adaptation effects have a relevant impact in the BOLD response and, in particular, we will see that information of the recurrent neuronal activity can be extracted from the post-stimulus undershoot of the BOLD signal.

### 3.2. Calcium coding and BOLD response

In this section we analyze how the information about the input stimulus and the neuronal activity is codified and transmitted by the calcium signal to the vascular system. In particular, we focus on how the intensity of the stimulus (represented by the external input *v*_*ext*_) is codified. We separate the coding of the input stimulus into two parts: the first coding performed by the neuronal activity and the subsequent coding of the neuronal activity performed by the calcium signal. In Fig.4.a we show the response of the excitatory neuronal population as a function of the external input *v*_*ext*_, obtained from the mean field model (Eqs. 1,2). The curve corresponds to the steady value of the neuronal activity, after *v*_*E*_ and *W* have reached an equilibrium. The response exhibits a non-linear behaviour with signs of saturation for the highest inputs. This neuronal activity is then sensed by the astrocyte and codified by means of the intracellular calcium signal. We notice that the neuronal-astrocyte communica-tion is made through glutamate release, which in our model operates linearly on the curve of Fig.4.a. In Fig.4.b we show the time course of the calcium signal for two different inputs (*v*_*ext*_=3 and 6 Hz) and in panels (c),(d) we show the evolution of the frequency (*f*_*Ca*_) and amplitude (*A*_*Ca*_) of the calcium signal as a function of the neuronal (excitatory) firing rate (*v*_*E*_). It can be seen that the neuronal activity is codified by the calcium signal in terms of both its frequency and amplitude. We observe that the signal exhibits a threshold for calcium-spike generation, under which no spike is observed. Over the threshold, both amplitude and frequency are monotonic increasing functions of the firing rate. For a detailed analysis of the calcium dynamics and the coding modes see Supp. Inf. and Ref.[18].

**Figure 4:**
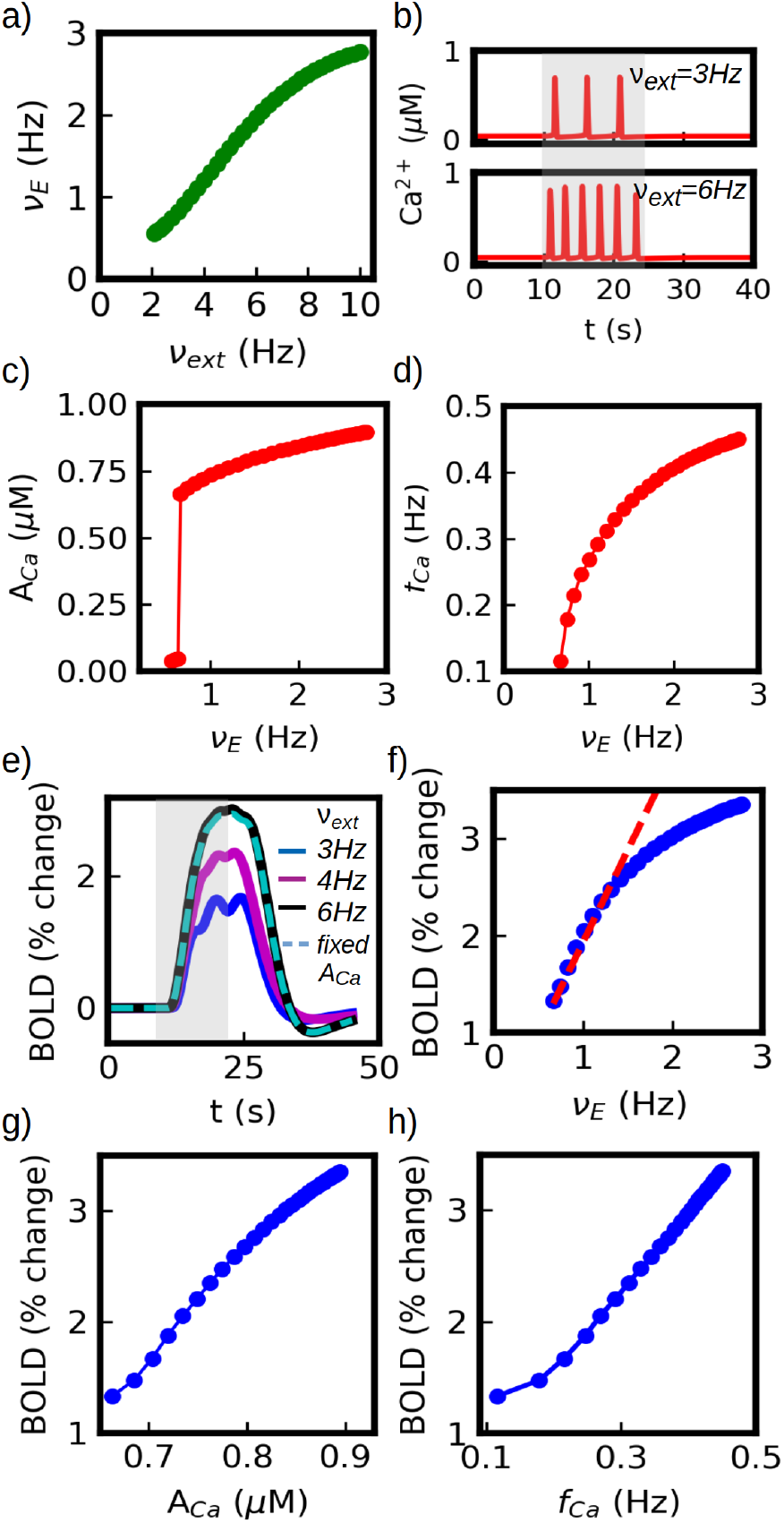
Neuronal and calcium coding of an external stimulus. a) Average firing rate of the excitatory neuronal population as a function of the external stimulus intensity (measured in terms of the external excitatory rate *v*_*ext*_). b) Astrocytic calcium activity for two values of *v*_*ext*_. c-d) Amplitude and frequency of calcium-spikes as a function of neuronal (excitatory) firing rate. External stimuli are first codified by neuronal activity which in turns modulates both the amplitude and frequency of the calcium response. e) BOLD response for different stimulus intensity (for a comparison with experimental results see Fig. 2.b). In dashed line we show the results for *v*_*ext*_=6Hz with the calcium-spike amplitude limited to A_*Ca*_=0.65 *μ*M (amplitude at threshold) for arachidonic acid production, indicating that the main increase in BOLD response with stimulus intensity is not driven by the change in calcium amplitude. f) BOLD response as a function of neuronal (excitatory) activity. A linear ear fit is shown in red dashed line, which describes ∼75% of the total BOLD variation. g-h) BOLD response to stimuli intensity parameterized in terms of calcium-spike amplitude and frequency. For a comparison of panel (g) with experimental results see Ref.[11].

Finally, the calcium signal is transmitted to and decodified by the vascular system through the arachidonic acid avalanche and cAMP pathway as explained in the previous sections. In Fig.4.e we show the BOLD response for three different inputs (*v*_*ext*_=3,4 and 6 Hz). We see that the variation in the input performs basically a re-scale in the amplitude of the BOLD signal.

In Fig.4.f we show the amplitude of the BOLD response as a function of the neuronal activity *v*_*E*_. A linear fit is shown in red dashed line, which describes nearly 75% of the total BOLD variation in our simulations. After this linear regime, a saturation in the response is observed. In Fig.4.g-h we show the BOLD response to input intensity parameterized in terms of calcium-spike amplitude and frequency. A similar relation between BOLD and the amplitude of astrocytic *Ca*^2+^ signal has been seen in experiments [11]. Nevertheless, we notice that the change in BOLD response in our simulations is mainly driven by the frequency of the calcium signal and not by its amplitude. We show this in Fig.4.e (dashed line) by imposing a fixed *A*_*Ca*_ for arachidonic acid production. The change in *f*_*Ca*_ accounts for more than 95% of the BOLD amplitude variation, while the other %5 is given by the change in *A*_*Ca*_. In addition, we see that the BOLD-*f*_*Ca*_ relation is approximately linear, except near the threshold where the linearity is lost.

The threshold for calcium-spike generation depends strongly on the maximum rates of glutamate-dependent IP3 production and on the rate of IP3-induced calcium realease from endoplasmic reticulum. Dysfunctions in the rate of calcium release (for example, by variation in the density of IP3 receptors in the membrane of the endoplasmic reticulum) may lead to an increase in the threshold level of glutamate concentration for calcium-spike activation, meaning that a higher synaptic activity would be necessary to evoke a significant hemodynamic response. Further increase in the threshold level would lead to a decoupling of the neurovascular response (an example of such a dysfunction is presented in SI).

### 3.3. Linear coupling and the Hemodynamic Response Function

The BOLD response shown in Fig.2.a for a single short (< 4s) stimulus is known as the Hemodynamic Response Function (HRF). In fMRI experiments it is usually assumed that the neuronal and vascular activity are linearly coupled, which has shown to be valid in a certain parameter range [6, 35]. Assuming that the coupling is linear and time invariant, it is possible to write the vascular (BOLD) response as [35]:

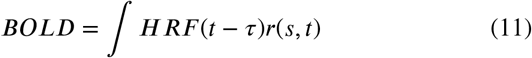

where *r*(*s, t*) is the time course of neuronal activity of the local population, being *s* the intensity of the stimulus or other stimulus parameter under study. In this section we study the HRF obtained from our model and the validity of the linear analysis. The HRF obtained for a stimulus of 2 seconds and *v*_*ext*_=4Hz is shown in detail in Fig.5.a. Main features of the HRF (indicated in the figure) observed in experiments comprise: i) a lag of *t*_*ON*_ ≈2 seconds between the application of the stimulus and the activation of the response, ii) a lag of *t*_*peak*_ ≈5 seconds between the application of the stimulus and the peak of the response, iii) a maximum amplitude that ranges between 1% and 2% (measured as percentage change from the basal level) and iv) a marked after-stimulus undershoot [35, 33]. As shown in Fig.5.a, all these features are correctly captured by our model. For completeness we show in the figure a fit performed with the so called canonical HRF, a phenomenological expression widely used for fMRI data analysis. As we see the HRF obtained from our model is in good agreement with the canonical HRF.

In Fig.5.b-c we show the BOLD response obtained from the simulations together with the results from Eq.11 (for a pulse and an oscillatory stimuli respectively). As we can see the HRF formalism can correctly reproduce the simulations result within the predicted linear range. Thus, in our model the calcium pathway generates a neurovascular coupling that behaves linearly close to the threshold of BOLD (and calcium) activation (*v*_*ext*_ ≈ 2.4) and for a range of more than a twofold change in neuronal activity which accounts for a about 75% of the total change in BOLD signal in our simulations.

Based on the analysis performed so far and the results shown in Fig.4, we propose now to use the calcium activity as a predictor of the BOLD response instead of the neuronal activity. In Fig.4.h we can see that the BOLD follows a linear relation with the calcium firing rate (*f*_*Ca*_), which makes it a good candidate for the HRF formalism. Thus, we will replace the function *r*(*s, t*) from Eq.11 by *c*(*s, t*), corresponding to the calcium activity. In Fig.5.d-e) we show the results of the convolution between HRF and the time course of the calcium activity. We see that the convolution can correctly reproduce the results of the simulations for barely the entire range, except close to the threshold where the BOLD-*f*_*Ca*_ relation deviates from the linearity.

**Figure 5:**
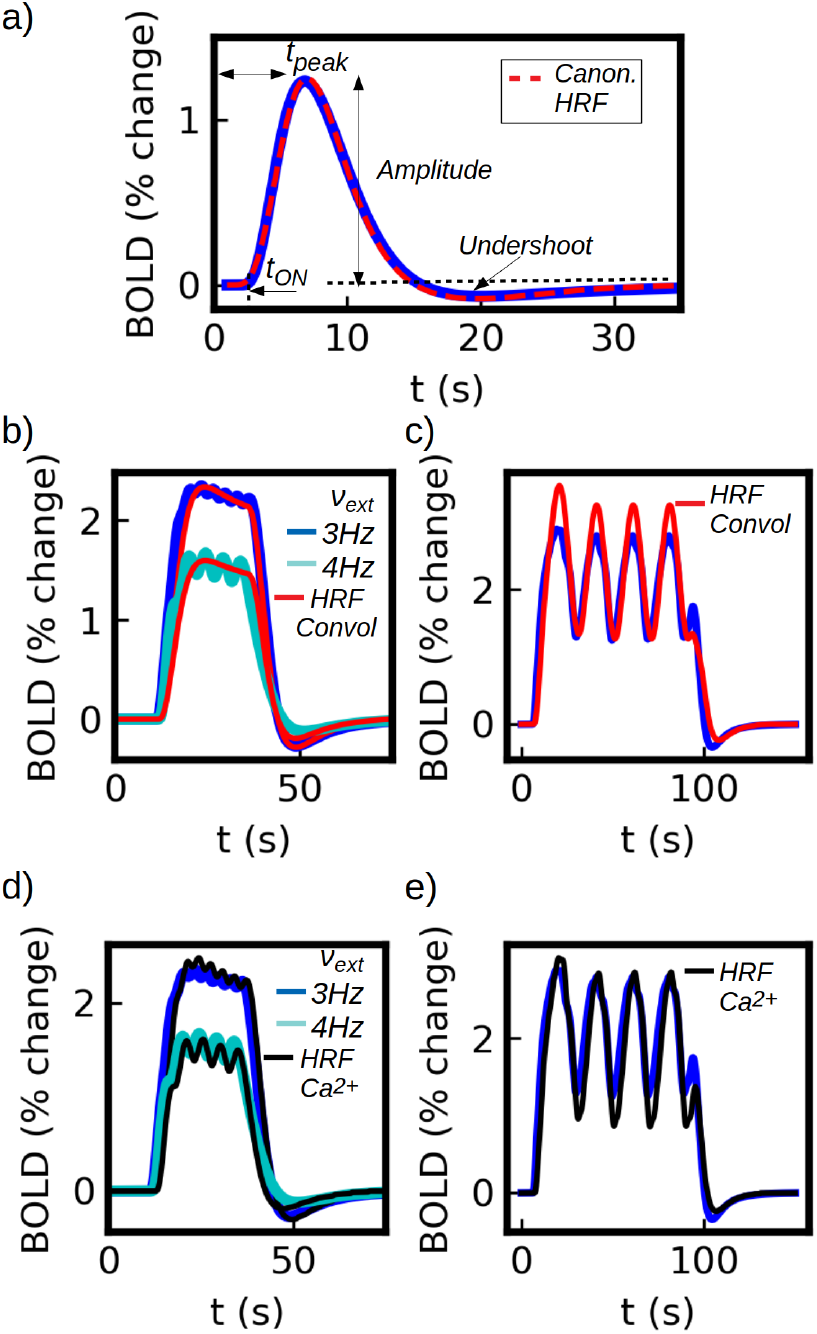
a) Hemodynamic Response Function (HRF) obtained from our model, corresponding to a stimulus of 2 seconds and *v*_*ext*_=4Hz (blue solid line). A fit with a double-gamma canonical HRF is shown (red dashed line), 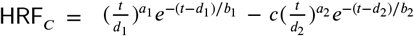. The coefficients from the fit are *d*_1_ = 6s, *a*_1_ = 6,*b*_1_ = 1*s,c* = 0.07, *d*_2_ = 18s, *a*_2_ = 12,*b*_2_ = 1.5s, in good agreement with experimental values [36, 37]. In panel (b) and (c) we show the BOLD response together with the result from the convolution of Eq.11. In panel (b) we show the results for a 24 seconds stimulus with *v*_*ext*_=3 and 4Hz. In panel (c) we show the results for a sinusoidal stimulus centered at *v*_*ext*_=4.2Hz and amplitude of 1.6Hz. We see that the results from the convolution fails to reproduce the simulations for the higher values of the input as the system moves away from the linear regime. In panel (d) and (e) we show the BOLD response estimated via the convolution of the HRF with the calcium activity (for the same stimuli as panels (b)-(c) respectively). From panel (e) we see that the results from the convolution can reproduce the simulations except for the lower values of the input (close the threshold for calcium activation) as the system moves away from the linear regime.

In this section we have proved that the calcium activity is a good indicator of the BOLD activity and that it can be incorporated to the linear HRF formalism, exhibiting a range of validity larger than its neuronal counterpart. In addition, the fact that the HRF formalism is compatible with our model (and that our HRF is equivalent to the canonical HRF), indicates that the analysis performed in this paper for the calcium pathway of neurovascular coupling can be generalized to other models. Independently of the exact path that links the calcium activity with the vascular system, we have shown that the calcium-vascular link can be replaced by the linear HRF formalism using a canonical HRF which are experimentally tested and widely accepted. This makes the analysis presented in the following sections extensible to other alternative paths as far the linearity of the calcium-vascular relation remains valid. The linear and nonlinear regimes of the coupling in our model depend mainly on the transfer function (frequency response) of the astrocytic calcium dynamics. The origin of the saturation can be traced to both the IP3 and the pure calcium dynamics from the Li-Rinzel model. In the Li-Rinzel model, the frequency of calcium-spikes is given by the concentration of IP3 and it exhibits a frequency-lock for sufficiently high concentrations of IP3 (i.e. the frequency remains unchanged for higher values of IP3). In addition, the IP3 dynamics used in our model (which depends on the glutamate concentration) exhibits a saturation in the maximum concentration of IP3 for sufficiently high values of glutamate. The glutamate and IP3 concentrations at which saturation appears depend primarily on the maximum rates of glutamate-dependent IP3 production and on the affinity of IP3-receptors in the membrane of the endoplasmic reticulum that mediate the calcium release. Finally, we notice that, although the HRF is driven by the calcium dynamics (being a single calcium-spike the minimum activation ‘unit’), the characteristic times of the HRF (*t* _*ON*_, *t* _*PEAK*_) depend on the whole dynamics of the system. Thus, even when the calcium-spike frequency changes with the stimulus intensity, these changes have a relative small impact in the characteristic times of the HRF. The fact that the shape of the HRF does not change with the stimulus strength is a fundamental feature for the validity of the HRF formalism (see, for example, time-contrast separability in ref.[35]).

### 3.4. BOLD undershoot and calcium dynamics

One characteristic feature of the BOLD response is the presence of a post-stimulus undershoot. The origin of this undershoot is usually attributed to three main sources: a) a slow recovery of the venule volume [38, 19]; b) sustained post-stimulus metabolic rate of oxygen [27]; and c) a post-stimulus undershoot in cerebral blood flow (CBF) [38]. The first of this sources is captured by the Balloon model [19] and is thus incorporated in our simulations. In this section we will focus on the third of this sources, namely the post-stimulus undershoot in CBF. The second source is not incorporated in the current version of our model, but it might be object of future analysis.

In our model the level of CBF is modulated by the calcium activity. In Fig.6.a we show the time course of the calcium signal and the CBF_*IN*_ during the application of a stimulus. We see that after each calcium-spike there is a period where the Ca^2+^ level goes below its basal state. This is generated by the fast decrease of IP3 levels driven by the calcium-spike (for details see Supp. Inf. Materials and Methods). While the stimulus is applied the IP3 levels rise again and a new spike is generated. However, in the post-stimulus phase the IP3 level slowly recovers until it reaches its basal concentration. During this last period the Ca^2+^ level also remains below its basal concentration giving place to a post-stimulus undershoot in the calcium signal.

**Figure 6:**
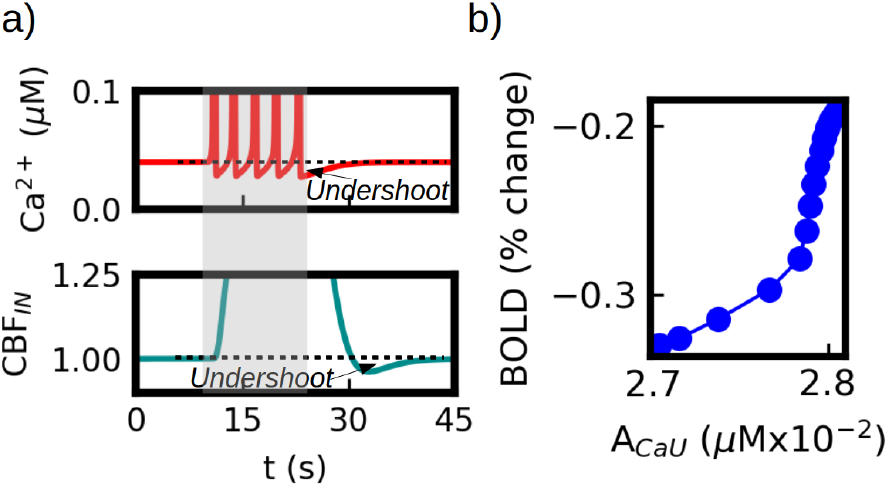
Calcium effects in BOLD post-stimulus undershoot. a) Top panel: calcium time course during stimulation. After the stimulus is removed the calcium signal remains below its basal level and slowly recovers (indicated as *Undershoot* in the figure). The depressed calcium level leads to a post-stimulus undershoot in the blood flow (CBF _*IN*_) which is shown in the bottom panel. b) Evolution of the BOLD undershoot amplitude as a function of the calcium undershoot amplitude. We see that the changes in the calcium level lead to a variation of 50% in the BOLD undershoot.

This undershoot is transmitted to the CBF_*IN*_ as shown in Fig.6.a (bottom panel). The size and duration of this undershoot depends on the intensity of the stimulus and on the phase of the calcium oscillation. In Fig.6.b we show the relation between the amplitude of the BOLD undershoot and the amplitude of the calcium undershoot (A_*CaU*_), measured as the minimum value of the BOLD and calcium signals respectively. We see that the calcium undershoot in our simulations can account for up to 50% change of the BOLD undershoot.

### 3.5. Adaptation Effects

In this section we analyze the effects of neuronal adaptation in the neurovascular coupling. In our model the adaptation is described within the mean-field formulation by the variable *W* (see Eqs. 1 and 2), which accounts for the average adaptation of the AdEx neuronal population. We focus here only on the spike-triggered adaptation characterized by the parameter *b* (we use *a*=0 in Eq.2). The action of adaptation in neuronal activity can be seen in the first and second panels (from the top) of Fig.3. At the onset of the stimulus the neuronal activity exhibits a large overshoot generated by the interplay of the fast neuronal response and the slower adaptation variable *W*. Similarly, an undershoot in the neuronal activity is observed after the end of the stimulus, generated by the same mechanism. This dynamics is also captured by the glutamate concentration, which follows the variations in neuronal activity. If the characteristic time of the neuronal adaptation (given by *t*_*w*_) is much smaller than the characteristic time of the calcium response, then the adaptation effects become negligible for the neurovascular coupling and no effects are observed in the BOLD signal. This has been the case for the simulations presented in the previous sections, where we adopted *t*_*w*_ =1s.

The situation changes as the characteristic time of adaptation becomes comparable to the time of the calcium response. In Fig.7 we show the results for the case *t*_*w*_=5s. In panel (a) we show the neuronal (top) and BOLD responses (bottom) and we compare them with results having no adaptation (*b*\*=0, see caption in Fig. 7). In this case the adaptation effects are clearly observed in the BOLD signal. At the onset of the stimulus we observe the larger BOLD response in the blue curve (*b* ≠ 0) driven by the initial neuronal overshoot. Although the variation is small, we see that the response is not only larger but its maximum value is reached earlier than expected when no adaptation is considered, which is a feature observed and discussed in experiments [35]. In addition, after the stimulus, we see a large variation in the amplitude of the undershoot on the BOLD signal. These two effects can be traced to the calcium dynamics shown in panel (b). Here we can see the effects of the adaptation acting on the frequency and amplitude of the calcium-spikes at the onset of the stimulus: the initial increase in glutamate concentration generated by the neuronal overshoot leads to a higher calcium frequency and amplitude which quickly decreases and reaches a lower steady state after a few seconds. This leads to the larger initial BOLD response. On the other hand, the post-stimulus neuronal undershoot generates a decrease in the post-stimulus calcium concentration which is the origin of the larger undershoot observed in BOLD response. In panel (c) we show the variations in the BOLD undershoot as a function of the adaptation parameter *b*. We see that the post-stimulus neuronal undershoot can account for up to 70% of the BOLD undershoot for the larger values of *b* (for the details of the plot see caption). We notice that this neuronal dynamics also leads to a dependence of the PSU on the duration and strength of the stimulus. For short stimulus (compared to the *W* response) the influence of neuronal adaptation will be limited, for which a smaller neuronal-driven PSU in the BOLD signal is obtained.

**Figure 7:**
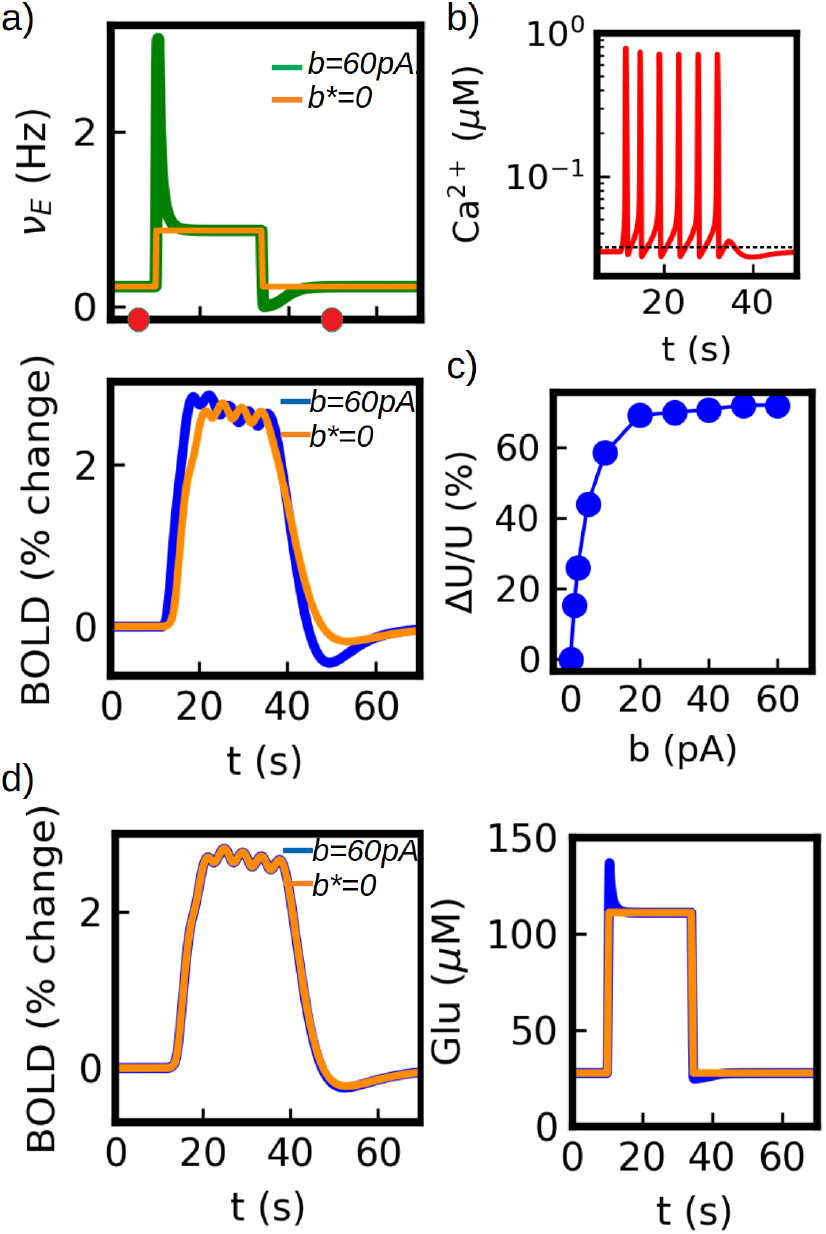
Neuronal adaptation effects in BOLD signal. a) Top: neuronal activity during stimulus application with (green) and without (orange) adaptation (for a better comparison, we rescale the asymptotic firing rate for *b* = 0 to match the one of *b* = 60pA, so we indicate it with *b*\*). Bottom: BOLD response with (blue) and without (orange) adaptation. The effects of adaptation can be observed in the higher initial response for the blue curve and in the larger post-stimulus undershoot. The simulations correspond to *t*_*w*_=5s. b) Calcium-spike adaptation. The frequency and amplitude of the calcium-spikes are modulated by the changes of glutamate concentration due to neuronal adaptation. In the figure we can see the adaptation effects occurring during the first three calcium-spikes after which a steady state is reached. In addition, a decrease in the post-stimulus calcium concentration is induced by the neuronal post-stimulus undershoot. The calcium time course shown correspond to the lapse within the red circles indicated in top panel (a). c) Adaptation effects in the post-stimulus undershoot (PSU) of the BOLD response. The plot shows the variation in the PSU of the BOLD response as a function of the adaptation constant *b* (see Eq. 2). The variations in the undershoot are measured as ΔU/U=(min(BOLD)-min(BOLD_*Ad*_))/min(BOLD), where min() indicates the minimum value and BOLD_*Ad*_ correspond to the BOLD response when neuronal post-stimulus undershoot is removed. In our simulations the undershoot of the neuronal activity is responsible of up to about 70% of the BOLD PSU for the highest values of *b* (with t_*W*_ =5s). d) Simulation results for *g*_*r*_ = *g*_*ext*_ (equal contribution of external and recurrent activity to glutamate release). We show the BOLD response (left) and glutamate concentration (right). In this case the effects of neuronal adaptation on the BOLD signal are neglectable. The comparison with previous panels (*g*_*ext*_ = 0), indicates that information about relative role of recurrent activity can be extracted from the post-stimulus undershoot.

Finally, we notice that the results presented so far correspond to the case *g*_*ext*_ = 0 in Eq.3, meaning that the only source of glutamate release is the recurrent synaptic activity (which represents the output of the local neuronal population). As we explained in previous sections, when *t*_*w*_is small then the results obtained for the BOLD signal are nearly independent of the choice of *g*_*r*/*ext*_. However, when *t*_*w*_ is increased and the adaptation effects are captured by the BOLD signal as in this section, then the relative contribution of the recurrent and incoming synaptic activity to glutamate release becomes relevant (where we assume that the incoming activity exhibits no adaptation). In Fig.7.d we show the results for the case *g*_*ext*_ = *g*_*r*_ (i.e. equal contribution). We show in the figure the time course of the glutamate concentration (right panel), where we can see that the effects of the neuronal adaptation are drastically reduced in comparison to the case with *g*_*ext*_ = 0. Furthermore, we can see that there is no observable variation in the BOLD signal (left panel) for the simulations with or without adaptation. In particular, the post-stimulus undershoot in the BOLD signal is no longer influenced by the neuronal adaptation. This indicates that the amplitude of the PSU depends on the relative size of the input and output (recurrent activity) of the local neuronal population. Thus, one possible interpretation is that the amplitude of the BOLD signal provides information mainly about the input to the neuronal population, while the amplitude of the PSU provides information about its output.

## Discussion

In this paper we have presented a new framework for modeling neurovascular coupling. This framework is centered on calcium activity in astrocytes, that acts to link neuronal activity to the vascular system. Starting from a relatively detailed description of the coupling we have focused on fundamental aspects of the fMRI phenomenology, such as the different experimental designs, the Hemodynamic Response Function (HRF), the linearity in the coupling, the post-stimulus undershoot and the effects of neuronal adaptation. We have shown that calcium signaling plays a relevant role in all of these processes. Previous papers with models of the neuro-astro-vascular interaction have been proposed [14, 15, 16], but to the best of our knowledge, the present work is the only one that accounts for the above experimental observations on fMRI.

We have shown that the calcium-driven HRF generated by our model is equivalent to the one observed experimentally and we provided a comparison with the canonical HRF.

We have seen that calcium signaling can explain the linear and non-linear features of the neurovascular coupling. The non-linear features involve the existence of a threshold of neuronal activity for vascular activation and saturation in the BOLD signal for sufficiently high neuronal activity. In parallel our simulations predict a dynamical range where the coupling is linear. The linear coupling comprises around 75% of the total variation in the BOLD signal explored in our simulations and corresponds to the typical range of values seen in experiments (between 1 and 3 %). We have shown that within this range we can reproduce our simulations with the HRF linear formalism, which is widely used for data-analysis of fMRI experiments. In addition our model predicts that the BOLD signal is linearly coupled with the calcium activity, for which we have tested the HRF formalism with the calcium activity (instead of neuronal) which provides better results.

The way that calcium signal codifies and transmits information from the neuronal to the vascular system was studied. We have found that, in our model, the coding is performed mainly via frequency modulation of calcium-spikes with a small contribution of amplitude modulation (see Supp. Inf. for a discussion of the BOLD response to neuronal activity at different frequencies).

Other important aspect studied in the paper is the role of calcium signal on the BOLD post-stimulus undershoot (PSU). We have explored the contributions of neuronal and calcium activity to the PSU. We have shown that an undershoot in calcium concentration, which emerges from the same calcium dynamics, has a relevant contribution to the PSU. The recovery of the basal calcium concentration occurs faster than the the recovery of the BOLD PSU, which indicates that calcium might have a stronger contribution on the early phase and on the amplitude of the PSU. Such a calcium undershoot can be observed experimentally [10, 39, 40, 41, 42] but, to the best of our knowledge, has not been studied so far in relation with the BOLD signal.

In addition, we have shown that the inclusion of neuronal adaptation generates a post-stimulus undershoot in the neuronal activity which is captured by the calcium signal (coded in terms of calcium-spike frequency and amplitude adaptation) and transmitted to the vascular PSU. The contributions of PSU described here operate on the BOLD via the CBF, i.e. they lead to an undershoot in the CBF that is then captured by the BOLD. The existence of such neuronal-CBF-BOLD undershoot can also be observed in experimental work [38, 43]. We note that the undershoot of neuronal activity is independent of the astrocyte-calcium pathway and its impact on the BOLD might be taken into account for direct neuron-vascular interaction. Furthermore, it is believed that the direct neuron-vascular pathway also involves calcium activity as a mediator [6, 8], for which the results obtained from our model might be generalized for alternative neurovascular pathways. The impact of neuronal adaptation on BOLD signal has proven to be of importance in fMRI experiments and it has led to the development of fMRI-adaptation, a technique that makes use of neuronal adaptation and neuronal specificity. In this context the adoption of the AdEx formalism constitutes an important step towards an accurate description of adaptation effects in the neurovascular coupling, representing a relevant improvement with respect to standard neuronal-mass and field models used for fMRI analysis [44].

Possible extensions of our study may involve the description of BOLD imaging to a larger scale (multiple voxels or brain regions) and the analysis of resting state fMRI. The development of realistic models of the neurovascular coupling is currently of great relevance for the analysis of clinical data [45, 46]. Furthermore, the mean-field formalism adopted here provides tools to estimate electrical brain signals (LFP, EcoG), currently under development. This will give the chance to analyze multimodal experiments, where BOLD imaging is combined with electrical measurements, which is a topic of great interest [46]. Furthermore, our framework can be combined with different neuronal models and has the potential of being incorporated in brain simulators such as The Virtual Brain [29, 47].

In addition, since astrocytes play a role in several neurodegenerative diseases [48, 49, 50], our modelling framework may be investigated to predict how astrocytic dysfunction in brain disorders and injury can result in detectable changes in neurovascular coupling and BOLD signal. For example, mutations in the *PLA2G6* gene, associated to Parkinson’s disease [51], can lead to a disrupt in the release of Arachidonic Acid from membrane phospholipids and a reduction in calcium responses [50, 52], which could be captured by our model (see Supp. Inf. for an example of astrocytic dysfunction).

Finally, our model may be used for inferring neuronal activity from the BOLD signal, by reversing the equations (see Supp. Inf. for an example). In principle, this could yield to estimates of the global glutamatergic activity, and thus estimates of the population activity of excitatory neurons. Such estimates should be compared to simultaneous measurements of population activity and BOLD signal. As BOLD-fMRI is one of the leading non-invasive research tools in neuroscience, finding new and better methods to infer neuronal or calcium activity constitutes a promising direction for future work. Such methods can be of great relevance for studies of human brain networks in health and disease.

## Acknowledgments

Research supported by the CNRS and the European Union (Human Brain Project H2020-785907, H2020-945539).

## Code Availability

The code used for the simulations presented in this paper is available under request and will be available online after publication.

## Supplementary Information

## 1. Materials and Methods

### 1.1. Neuronal model

In our paper we consider a network made of Adaptive Exponential Integrate and Fire neurons (AdEx) [1]. The equations for the AdEx model are given by:

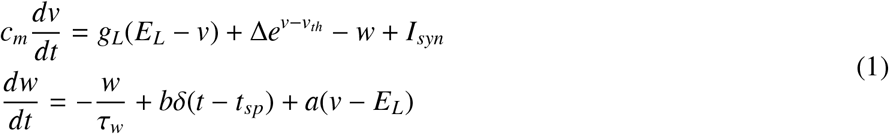

where *c*_*m*_ = 200 pF is the membrane capacity, *v* is the voltage of the neuron, and whenever v>v_*t*_*h* = 50 mV at time tsp(k), v is reset to the resting voltage *v*_*rest*_ = 65 mV and fixed to that value for a refractory time *T*_*ref*_ = 5 ms. The leak conductance is *g*_*L*_ = 10ns and the leakage reversal potential is *E*_*L*_=70 mV. The exponential term has a different strength for excitatory and inhibitory cells Δ = 2 mV (= 0.5 mV) for excitatory (inhibitory) cells. We consider inhibitory neurons with no adaptation (*a* = *b* = 0) and for excitatory neurons we take *b* = 60pA and *a* = 0. For the recovery time of adaptation we take *t*_*w*_ = 1s except otherwise indicated. The synaptic current *I*_*syn*_ received by a neuron i is the result of the spiking activity of all presynaptic neurons j ∈ pre(i) of neuron i. This current can be decomposed in the input received from excitatory E and inhibitory I presynaptic spikes 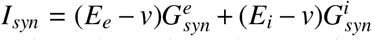, where Ee = 0 (Ei = 80 mV) is the excitatory (inhibitory) reversal potential. We consider voltage dependent conductances. We model 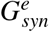 as a decaying exponential function that takes kicks of amount *Q*_*E*_ at each presynaptic spike,

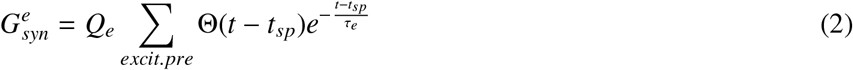

where Θ is the heaviside function, *τ*_*e*_ = *τ*_*i*_ = 5 ms is the decay timescale of excitatory and inhibitory synapses, and *Q*_*e*_ = 1.5 nS (*Q*_*i*_ = 5 nS) the excitatory (inhibitory) quantal conductance. We use the same equation with *e*→ *i* for inhibitory neurons.

The use of AdEx neurons provides a realistic description of neuornal activity and allows us to explore adaptation effects over the hemodynamic response. In addition, the spatial scale of the hemodynamic response suggests that vascular signals are affected by the population activity rather than by single neuronal dynamics. The strong correlation of the BOLD response with signals such as LFP also points in this direction [2]. Furthermore, each astrocyte in the brain interacts with thousands to millions of synapses [3, 4], suggesting that these cells are capable of sensing population activity. Thus, the choice of a mean-field formulation emerges naturally in this context.

### 1.2. Astrocytic Calcium Dynamics

To describe the astrocytic calcium dynamics we adopt a recent version of the Li-Rinzel model by De Pittá et al. [5, 6]. This model describes the activation of glutamate receptors in the astrocyte, the production and degradation of inositol trisphosphate (IP3) and the flux of *Ca*^2+^ between the cytosol and the ER of the cell. The fraction of activated glutamate receptors (*γ*_*A*_) is given by:

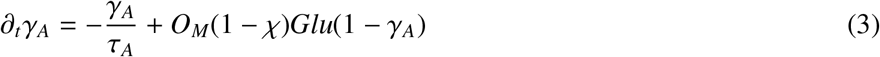

where (1 − *χ*) is the fraction of glutamate that spilled out of the synaptic cleft, *τ*_*A*_ is the characteristic receptor deactivation (unbinding) time constant, *O*_*M*_ is the binding rate and *Glu* is the glutamate concentration in the synaptic cleft. The values of the parameters are shown in Table 2.

**Table 2:**
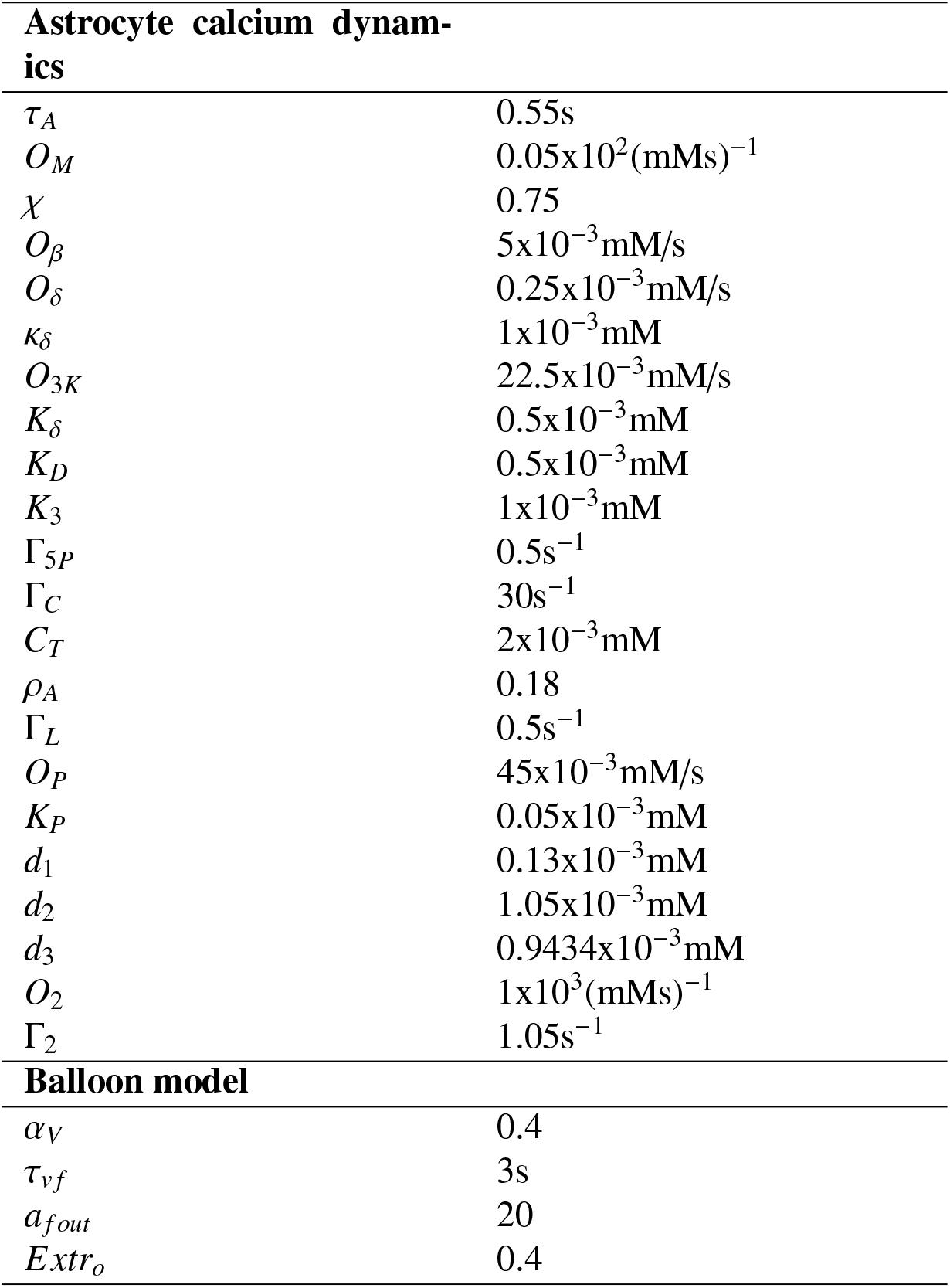
Model parameters for equations described in the SI

The IP3 concentration results from the *Ca*^2+^-modulated interplay of phospholipase C*β*-and C*δ*-mediated production and degradation by IP3 3-kinase (3K) and inositol polyphosphatase 5-phosphatase and evolves according to [6]:

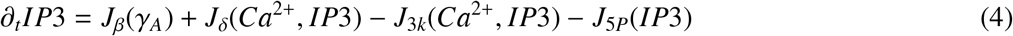

where

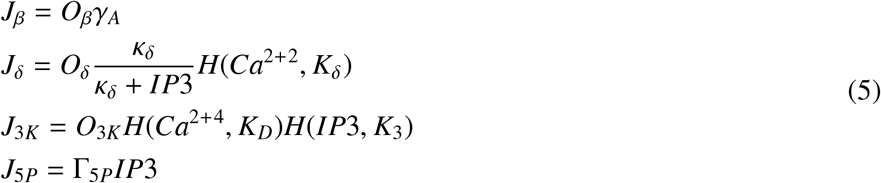

Here *H*(*x*^*n*^, *K*) denotes the Hill function *x*^*n*^/(*x*^*n*^+*K*^*n*^). Cytosolic calcium concentration and IP3 gating are described according to the Li-Rinzel model [7]:

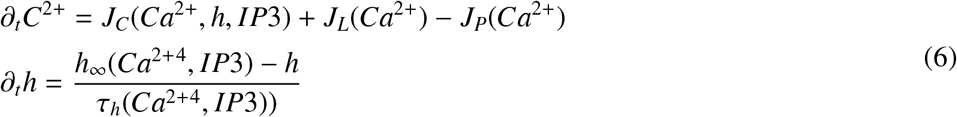

where *J*_*C*_, *J*_*L*_, and *J*_*P*_, respectively, denote the IP3-mediated *Ca*^2+^ -induced *Ca*^2+^ -release from the ER (*J*_*C*_), the *Ca*^2+^ leak from the ER (*J*_*L*_), and the *Ca*^2+^ uptake from the cytosol back to the ER by serca-ER *Ca*^2+^ /ATPase pumps (*J*_*P*_) [141]. These terms, together with the IP3R deinactivation time constant (τ_*h*_) and steady-state probability (*h*_∞_), are given by

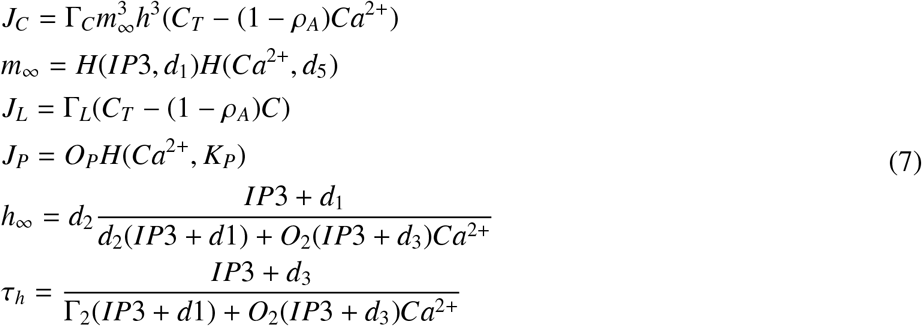

The values of the parameters are given in Table 2.

In this model the information about the input (i.e. glutamate concentration) is codified in terms of both the Amplitude and Frequency of calcium oscillations [5], as described in the main text. The relative relevance and efficiency of each type of coding depends on the set of parameters used and, in particular, in the duty-ratio of the calcium oscillations (i.e. the ratio between the width of the calcium pulse (spike) and the period) [5, 8]. Experimental and theoretical data suggests that the dominating coding mode of information is via frequency modulation [5, 9, 10, 11]. For this reason, we considered in this work relative low duty-ratio (<0.5) calcium oscillations which are more likely associated with efficient frequency coding [8]. On the other hand, high duty-ratio (smooth) calcium oscillations have been associated with the coordination of information between multiple astrocytes via intercellular signaling [5, 12]. Further analysis on the relative relevance of the coding modes can be an interesting topic for future studies, with the possibility that astrocytes may change the relation between coding modes depending on brain states, which could be of relevance for the analysis of resting-state fMRI.

Finally, we notice that the post-stimulus undershoot proper to the calcium dynamics studied in the main text depends on the IP3 oscillations around the basal state, which is a particular feature of the modifications introduced by De Pittá et al [5] to the original Li-Renzel model [7]. On the contrary, the post-stimulus undershoot driven by neuronal adaptation effects is the result of a decrease in the glutamate level (which in turns generate a decrease in the IP3 levels), and it is not specific to this particular model.

### 1.3. The Balloon model

To describe the generation of the BOLD signal we adopt the Balloon model [13, 14]. This model describes the change in dexohemoglobin concentration (which originates the BOLD signal) generated by an increase in CBF. The equations of the model are given by:

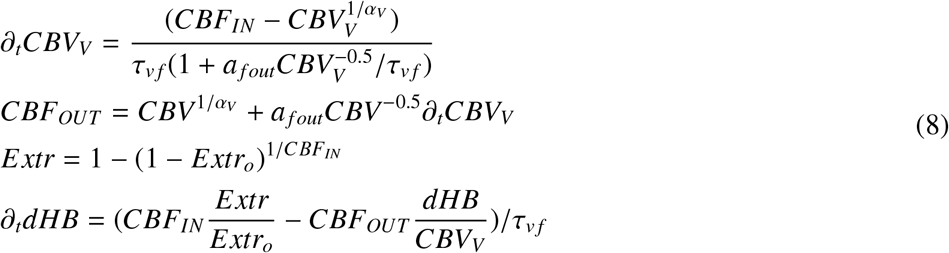

where *CBV*_*V*_ is the venous balloon volume, *CBF*_*OUT*_ is the flow of blood leaving the balloon, *Extr* is the oxygen extraction rate, *dHB* is the dexohemoglobin concentration. and τ_*vf*_ is the mean transit time through the venous compartment at rest.

### 1.4. Calcium-neurovascular pathway

We present here the simulations of the entire pathway for neurovascular coupling in our model. We show the results in Fig.1 and 2. In Fig.1 we show the main variables of the neuronal and vascular systems together with the calcium activity. In Fig.2 we show the the variables related with the arachidonic acid cascade and the cAMP dynamics.

In this paper we have focused on the excitatory synaptic activity as the main driver of the BOLD response. We notice that activity from inhibitory neurons has also been suggested as a driver for the BOLD response. It has been usually thought that excitatory synapses are the main drivers of the hemodynamic response as they represent the larger portion of energy consumption [15]. The inclusion of inhibitory neurons as a source of hemodynamics response in our model would be an interesting point for future studies. Nevertheless, we notice that excitatory and inhibitory activity are strongly correlated during stimulation [15]. In addition, the coupling with inhibitory neurons is believed to be carried either through astrocyte activation (similar to the description of our model) or by direct Nitric Oxide release which is also driven by intracellular calcium activity [16, 17]. Thus, we believe that the inclusion of inhibitory neurons as drivers of the hemodynamic response wouldn’t affect the main results of the current study.

**Figure 1:**
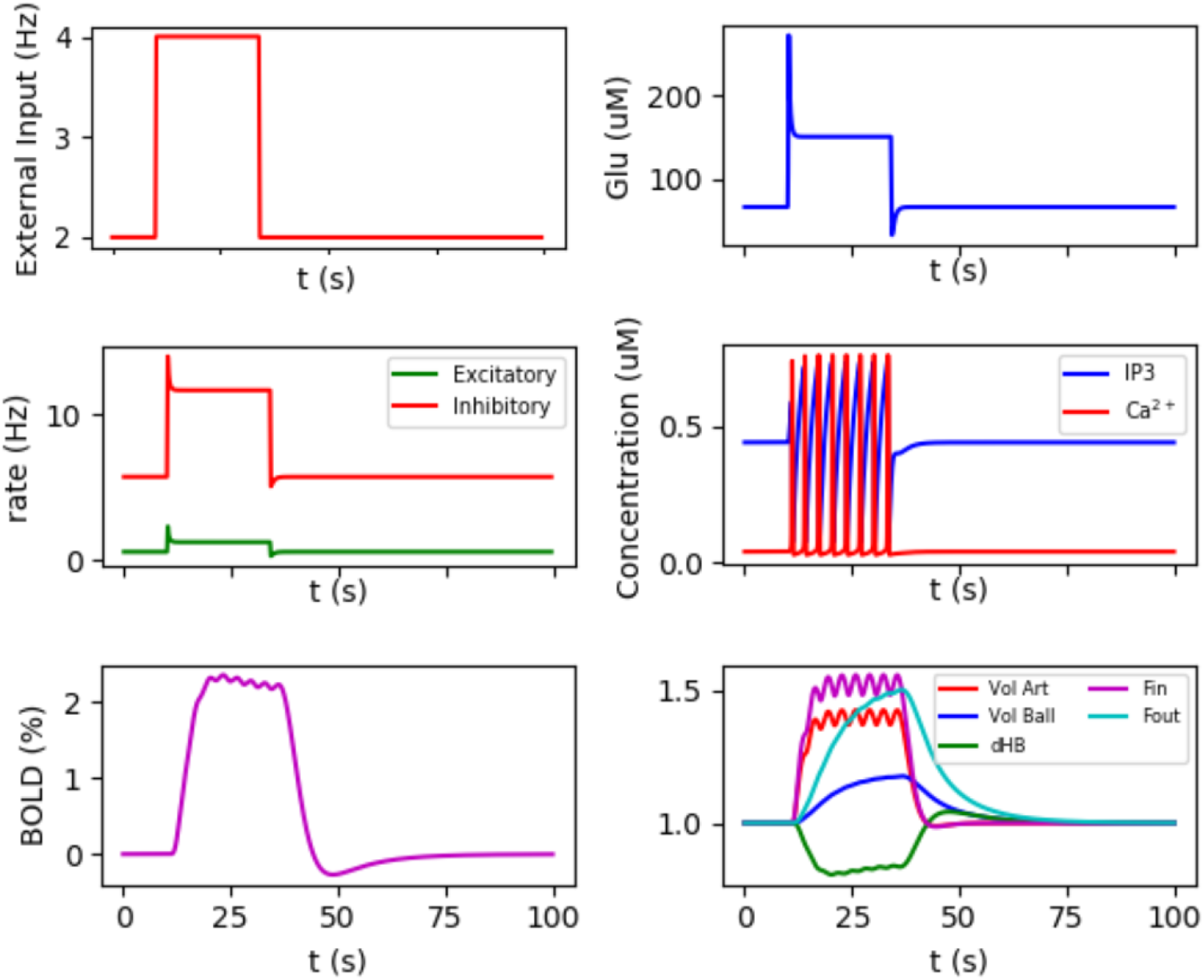
In this plot we show the main variables of the neuronal and vascular systems together with the calcium activity for a pulse of ν_*ext*_ = 4Hz and a duration of 24s.

**Figure 2:**
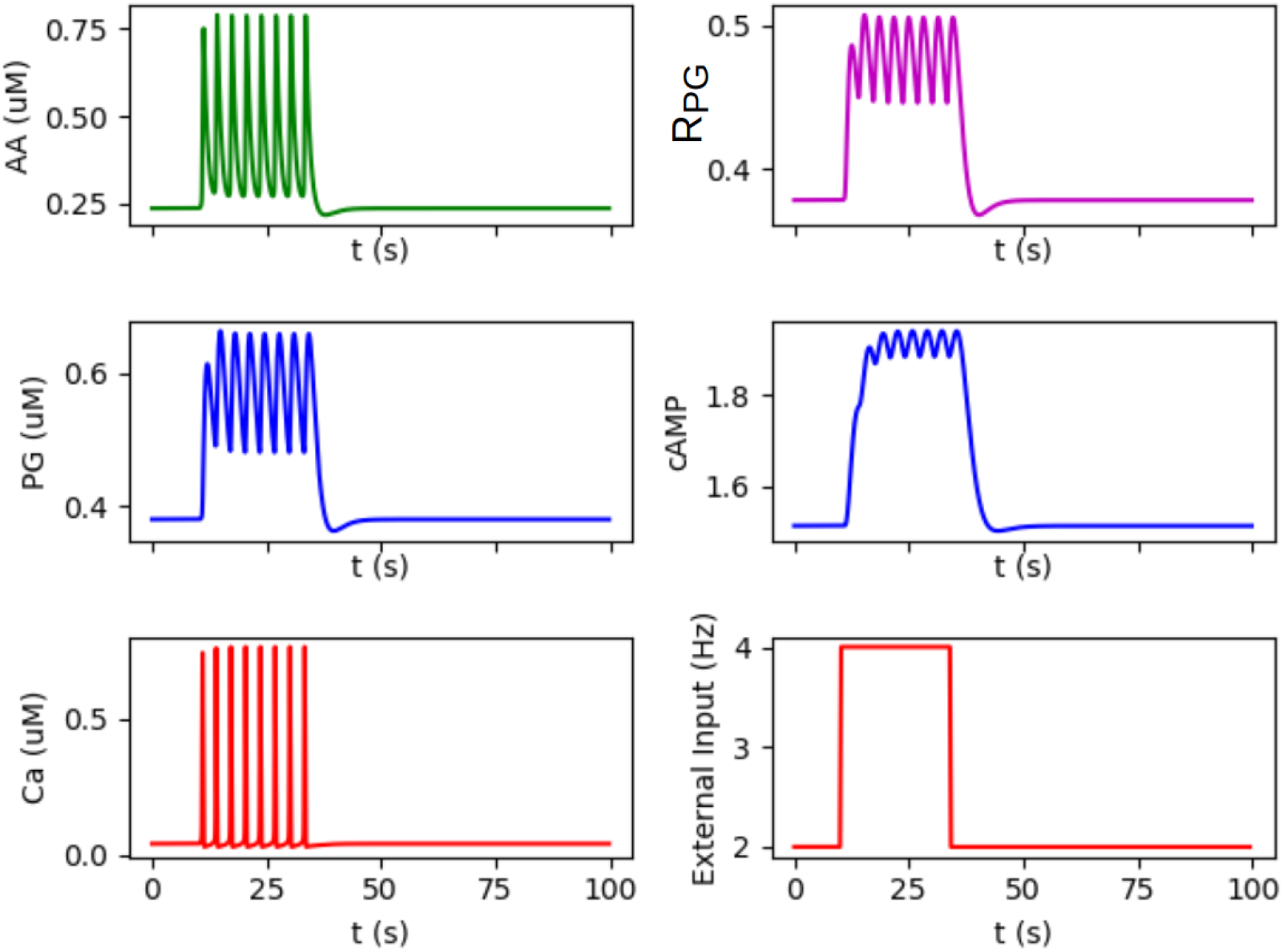
In this plot we show the variables related with the arachidonic acid cascade and the cAMP dynamics for a pulse of ν_*ext*_ = 4Hz and a duration of 24s.

## 2. BOLD response to neuronal activity at different frequencies

It is known that the BOLD signal exhibits a stronger correlation with Local Field Potentials (LFP) than with multi-unit neuronal activity (MUA) [2]. LFP is a signal related to the synaptic activity at the population level while the MUA reflects the activity of a small group of neurons (in the order of the ten(s)). Within the frequency range of LFP (0-100Hz) the correlation with BOLD has shown to be stronger for the gamma band (40-100Hz) [2, 18]. The preference for the gamma frequency has been associated to its stimulus dependent response, while frequencies within the beta band (18-30Hz) have shown to be stimulus-independent and may reflect the contribution of a stimulus-independent neuromodulatory pathway [18, 19]. In addition, the alpha band (8-12Hz) has been seen to be anti-correlated with the BOLD signal (for constant total LFP spectrum power), which is attributed to the shift of the spectrum towards higher frequencies during the application of a stimulus [18]. We first notice that our model reflects the same higher correlation between the BOLD signal and the synaptic activity that is contained within the LFP. The neuronal activity in our model is described via a mean-field model which, by construction, describes the activity at the population level and not at the MUA level. In addition, the main source of LFP is known to be synaptic activity (rather than spiking), which in our model is represented by glutamate release and is the driving signal for the calcium dynamics in astrocytes and the consequent BOLD response. Thus, in our model the BOLD response reflects the synaptic activity at the population level in equivalence to the LFP. Second, in our model the BOLD is stimulus-dependent for which a higher correlation with the stimulus-dependent LFP band (gamma) would also be expected. Third, the shift of the spectrum toward higher frequencies is also implicit in the mean-field model via the variation of the mean firing rate during stimulation. Thus, our model is in qualitatively agreement with the experimental observation regarding the dependence of the BOLD with the LFP signal and its frequency components.

## 3. Astrocyte dysfunction and neurovascular coupling

In this section we present an example of a dysfunction in astrocytes in our model that has repercussions in the neurovascular coupling. In Fig.3 we present the calcium and BOLD response when the maximum rate of calcium release from the ER towards the cytosol (Γ_*C*_) is altered (see Eq.7). We see how this alteration generates first a reduction in the frequency of the calcium spikes and, as the alteration is increased, the spiking dynamics is completely suppressed. The alteration is also observed in the BOLD response (panel (b)), where the amplitude of the signal is reduced. In panel (c) we show the power spectrum of the BOLD response. The peak in the spectrum corresponding to the frequency of calcium spikes can be detected in the BOLD signal. This peak exhibits a shift toward lower frequencies for the altered astrocyte (middle panel) and the two first harmonics can also be detected.

**Figure 3:**
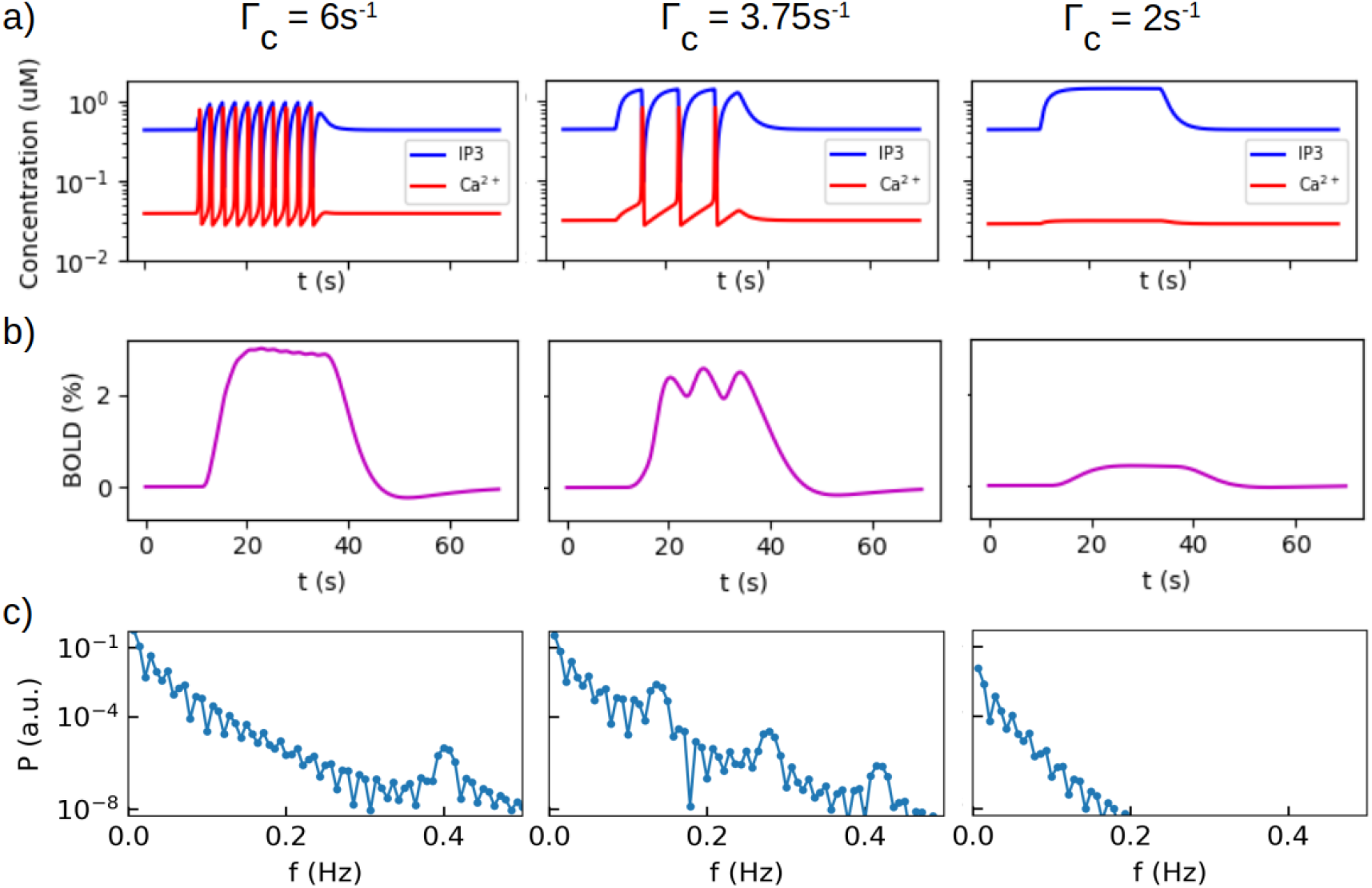
Astrocyte dysfunction and neurovascular coupling. We show simulations for three different values of maximum rate of calcium release from the ER towards the cytosol Γ_*C*_ (see Eq.7). a-b) Calcium dynamics and BOLD response for a simulations of ν_*ext*_ = 6*Hz* applied for 24s. c) Power spectrum of the BOLD signal. We see that the alteration in the astrocyte causes a reduction in the frequency of calcium spikes which induces a reduce response in the BOLD signal. In addition, the variation in frequency can detected in the power spectrum where the peak corresponding to the calcium dynamics exhibits a shift toward lower frequencies, from 0.4Hz in the left panel to ~ 0.15Hz in the middle panel. The first two harmonics can also be detected in the later case. To compute the power spectrum the stimulation time was extended to 48s in order to obtain a significant number of spikes.

## 4. Neuronal activity estimation from the BOLD signal

The final goal of fMRI imaging is to obtain an estimate of the underlying neuronal activity. In this context realistic models of the neurovascular coupling represent a valuable tool. The model presented in this paper provides several paths to perform this task. The most direct method consists in a reversion of the neurovascular pathway via the spectral analysis of the BOLD signal. As seen in Fig.3.c, the frequency of calcium spikes can be detected in the spectrum of the BOLD signal (peak at 0.4Hz, left panel). Then, the excitatory neuronal activity can be directly obtained via the *f*_*Ca*_− ν_*e*_ relation (Fig.4.d, main text). In the case of Fig.3 (left panels a,b,c), the peak at 0.4Hz in the BOLD spectrum corresponds to ν_*e*_ = 2*Hz* which in turns correspond to *nu*_*ext*_ = 6*Hz* (see Fig.4.a, main text). The amplitude of the BOLD signal can also provide a similar estimation, although this is less direct and can be affected by different elements in the coupling pathway (i.e. variations in arachidonic acid response or even changes in the basal state). Further information about the neuronal activity can be obtained from the post-stimulus undershoot (PSU) in the BOLD response. As shown in the main text, the size of the PSU depends on the relative contribution to glutamate release of the recurrent (ν_*e*_) and incoming (ν_*ext*_) synaptic activity. In our model the contribution of ν_*e*_ and ν_*ext*_ is weighted by the proportionality constants *g*_*r*/*ext*_ which we use as free parameters in the main text. However, for analyzing real data, *g*_*r*/*ext*_ are given by the proportion of recurrent and incoming synapses in the network, which is a known value for some brain regions (for example 70/30% in the visual cortex). Thus, in this case ν_*e*_ and ν_*ext*_ can be estimated from the data. Furthermore, the mean-field model provides straightforward ways of calculating other brain signals such as Local Field Potentials, for which analysis of multimodal data can also be developed. This last is indeed a topic of great interest [20] which opens exciting paths of research for the future.

## Notes

### Competing Interest Statement

The authors have declared no competing interest.

